# A Model for the Peak-Interval Task Based on Neural Oscillation-Delimited States

**DOI:** 10.1101/448688

**Authors:** Thiago T. Varella, Marcelo Bussotti Reyes, Marcelo S. Caetano, Raphael Y. de Camargo

## Abstract

Specific mechanisms underlying how the brain keeps track of time are largely unknown. Several existing computational models of timing reproduce behavioral results obtained with experimental psychophysical tasks, but only a few tackle the underlying biological mechanisms, such as the synchronized neural activity that occurs throughout brain areas. In this paper, we introduce a model for the peak-interval task based on neuronal network properties. We consider that Local Field Potential (LFP) oscillation cycles specify a sequence of states, represented as neuronal ensembles. Repeated presentation of time intervals during training reinforces the connections of specific ensembles to downstream networks. Later, during the peak-interval procedure, these downstream networks are reactivated by previously experienced neuronal ensembles, triggering actions at the learned time intervals. The model reproduces experimental response patterns from individual rats in the peak-interval procedure, satisfying relevant properties such as the Weber law. Finally, we provide a biological interpretation of the parameters of the model.

## 1 Introduction

How the brain represents the passage of time and uses this information in time-related tasks is still an ongoing debate in the scientific community. There seems to be different mechanisms involved in temporal processing depending on the time scale (Buhusi and Meck, 2005; Paton and Buonomano, 2018). For example, interaural delay is key to temporal discrimination in the microseconds range (Shaffer, 1984), while circadian rhythms (Czeisler et al., 1999) seem to play a role in the estimation of many hours/days, besides controlling other important human functions such as body temperature (Benloucif et al., 2005). In particular, intermediate durations ranging from seconds to minutes (referred to as *interval timing* by the timing community) stand out in many aspects of animal behavior and physiology. Examples include foraging, decision making, sequential motor performance, and associative conditioning (Merchant and Lafuente, 2014; Buhusi and Meck, 2005).

There are several models of interval timing which are able to reproduce behavioral results. However, only a few can also describe the critical properties of timing observed in experimental data, such as proportional timing (i.e., the linear relationship between response latency and the interval timed), the scalar property (i.e., the linear relationship between the standard deviation of the response latency and the interval timed), and consequently the Weber’s law (the constant ratio between the standard deviation and the mean latency of the responses across different timed intervals) (Church and Meck, 2003). One of the most influential models which successfully accounts for these properties is the Scalar Expectancy Theory, SET (Gibbon, 1977; Grondin, 2014). SET is a cognitive process model which assumes the existence of a pacemaker which emits pulses at a certain rate that are stored by an accumulator. The temporal estimation comes from counting the number of pulses in the accumulator and comparing it to a number of pulses previously stored in the reference memory (i.e., past experiences). One common criticism to the model, however, is its lack of biological validity, as no pacemakers or accumulators have been identified in the brain.

Alternative models which do not necessarily assume cognitive intervening variables have been proposed, such as the Behavioral Theory of Timing (BeT) (Killeen and Fetterman, 1988) and a modified version of BeT called Learning to Time (LeT) (Machado, 1997; Machado et al., 2016). LeT, for example, assumes that different behavioral states are serially activated when an organism is experiencing the passage of time. Each of these states is linked to a response by some degree, and the strength of these associations (vector of weights) are updated at every reinforcement (in reinforced trials) and by its omission (extinction trials) during learning. Although LeT can reproduce many essential features of timing behavior such as the scalar property, it is subject to the same criticisms attributed to SET, that is, it lacks direct biological correlates for its main underlying assumptions.

In order to deal with the issue of biological validity, some biologically-inspired candidate models of timing have been proposed. One example is the Striatal Beat Frequency (SBF) model (Matell and Meck, 2004; Oprisan and Buhusi, 2011), where the coincidental activation of oscillating neurons underlies the mechanism of time-tracking. Indeed, natural brain oscillations play a crucial role in many aspects of behavior in the hippocampus (Klimesch, 1999; Belluscio et al., 2012), the thalamus (Steriade et al., 1993), striatum (Berke et al., 2004), amygdala (Halgren et al., 1977), and other areas. Interestingly, following the idea of series of active "states" proposed by LeT, areas such as the hippocampus are important for timing behavior (Meck et al., 1984) and appear to operate on sequences of states (Lisman et al., 2005; Buzsáki and Tingley, 2018), which could be chained using heteroassociative rules (Sompolinsky and Kanter, 1986; Camargo et al., 2018). The hippocampus has neurons, called *time cells* (Eichenbaum, 2014), that fire at specific moments during the timed interval. Studies have reported the existence of these cells in the medial entorhinal cortex (MEC) (Tsao et al., 2018) and in the medial prefrontal cortex (mPFC) (Tiganj et al., 2017).

Recent evidence also suggest that the dynamics of neural populations can account for behavioral data observed in timing tasks (Paton and Buonomano, 2018; Balci and Simen, 2016; Eichenbaum, 2014). In state-dependent networks (Karmarkar and Buonomano, 2007), for example, a downstream network can use the evolution of the neural state trajectory to measure the passage of time. In such models, the state of a network is defined as the set of active neurons at a given moment, in a clear parallel to the behavioral states proposed by LeT. Recently, it has been hypothesized that time is not explicitly represented in the brain but is a byproduct of ongoing activity which is composed of a succession of events (Buzsáki and Llinas, 2017). This dynamical arrangement also agrees with the idea of using a chain of events to perform time-related tasks. Although these biologically-inspired models can reproduce neural activity which is frequently observed, including those from ramping neurons (Durstewitz and Deco, 2008; Machens et al., 2005), only recently a recurrent neural network has been described which could satisfy Weber’s Law (Hardy and Buonomano, 2018).

In this article, we describe a candidate timing model inspired on brain networks with oscillatory activity and sequential state activation, in which states are represented as neuronal ensembles delimited by oscillatory cycles. We use the model to simulate data in a peak-interval procedure, where repeated presentations of time intervals during training reinforce the connections of specific ensembles to a down-stream network. Then, during a reproduction phase, these downstream networks are reactivated by previously experienced neuronal ensembles, triggering actions at the learned time intervals. We compare our simulations to real data collected from rats. The implementation of this model follows the proposal by LeT, providing a possible biological interpretation for its key assumptions.

## 2 Methods

### 2.1 Peak-Interval Procedure

We collected the data using the Peak-Interval Procedure, as shown in Figure 1. We trained six male Sprague Dawley rats initially to press a lever in a standard operant chamber to receive a food pellet (fixed-ratio 1 schedule of reinforcement for one experimental session). Next, we trained the rats to make a nose-poke response 20 seconds after the lever press in order to receive the food pellet (*tandem* fixed ratio 1/fixed interval 20 s schedule of reinforcement). During this phase, each trial initiated by the onset of a houselight. The first lever press after light onset initiated a 20-s fixed interval. Food was primed 20 s after the first lever press. The first head entry into the food cup (detected by the breaking of a photobeam) after food prime delivered the food pellet, terminated the houselight, and introduced a variable inter-trial interval (ITI = 40 s). If no head entries were made within 20 s from food prime, the houselight turned off, we initiated the ITI, but no food was delivered. We trained the rats for 98 experimental sessions in this phase to ensure behavior stability. Finally, in the last training phase, we introduced peak trials with a probability of 0.5, during which we recorded the responses (lever presses and head entries), but there was no food pellets. After 60 s from food prime, we terminated the houselight, introduced the ITI, and started the next trial. We just analyzed the peak trials from the last 15 sessions for this study. All experimental procedures were approved by the Institutional Animal Care and Use Committee at Brown University and conform to guidelines for the Ethical Treatment of Animals (National Institutes of Health).

**Figure 1:**
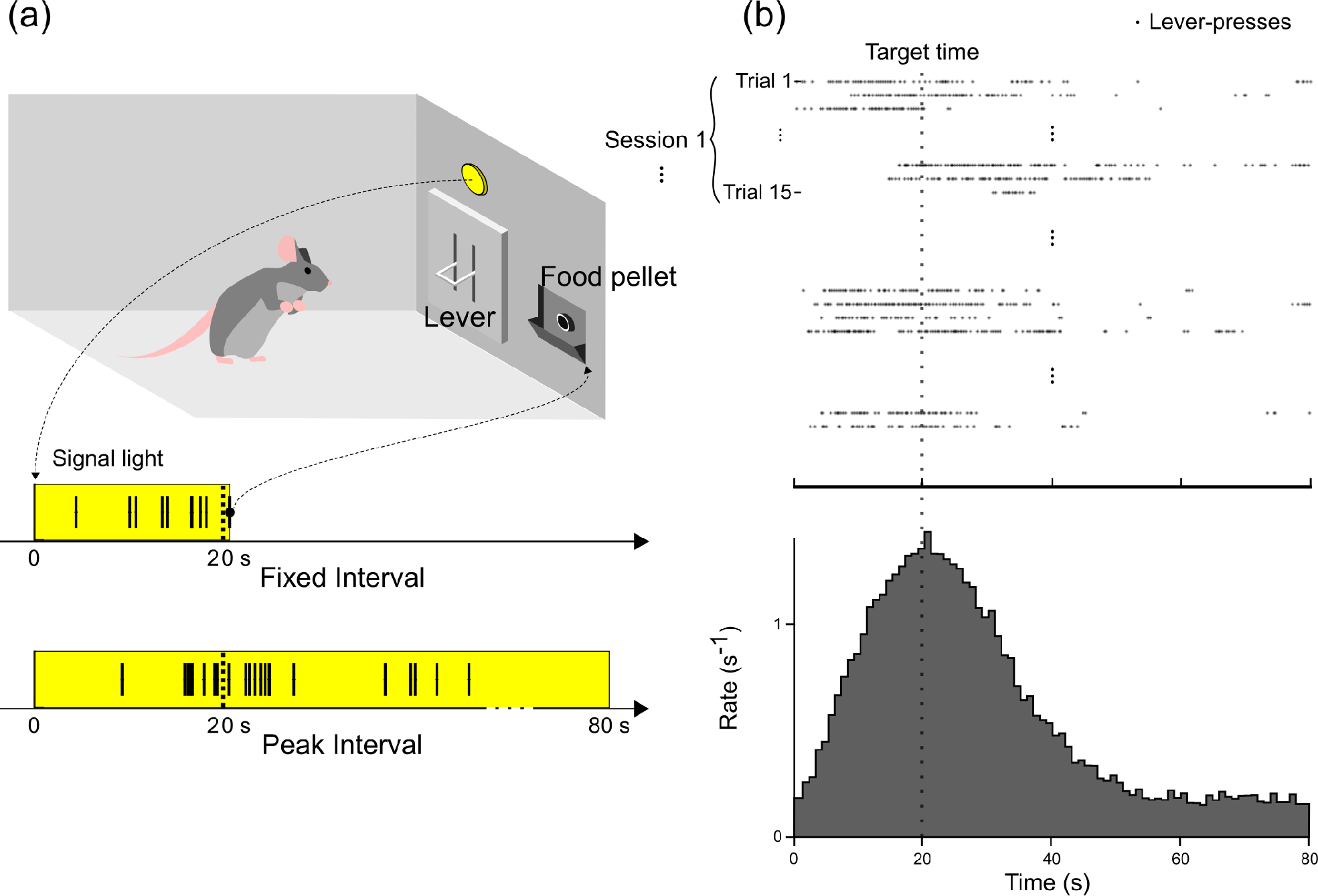
(a) Experimental setup of a typical Peak-Interval Procedure during a Fixed-Interval trial. A time-dependent light will be used as a stimulus and at the first lever press after 20 seconds the light will turn off and the food will drop at the food pellet. During a peak-interval, the light will keep turned on and the food will not drop. (b) Summary of the presses registered during each peak-interval trial with a target time of 20 s and the distribution containing the mean number of presses during each second of a trial.

### 2.2 Learning Mechanism

The model consists of a heteroassociative network represented by a series of successive states (Sompolinsky and Kanter, 1986; Camargo et al., 2018). Each state corresponds to a subset of active neural units within a Local Field Potential (LFP) oscillation. Each state *i* has a synaptic weight *w*_*i*_ connecting it to a downstream network. The weights store information about the specific time being learned, causing the downstream network to fire at that specific time. The model has two phases to account for the resetting of the downstream network for different time tasks: the learning phase and the reproduction phase.

During the learning phase, states are consecutively activated until a specific target time *T* has elapsed (Figure 2a). At that moment, the synaptic weight *w*_*i*_ between the current active state *i* and the down-stream network increases one (arbitrary) unit. This state activation sequence happens multiple times, one for each learning trial. For each trial, there is a transition time step *t*_*s*_ between states drew from a Gaussian distribution with mean *μ* and standard deviation *σ*. At the end of the simulation, we normalized the *w*_*i*_ values so that the maximum weight is 1.

**Figure 2:**
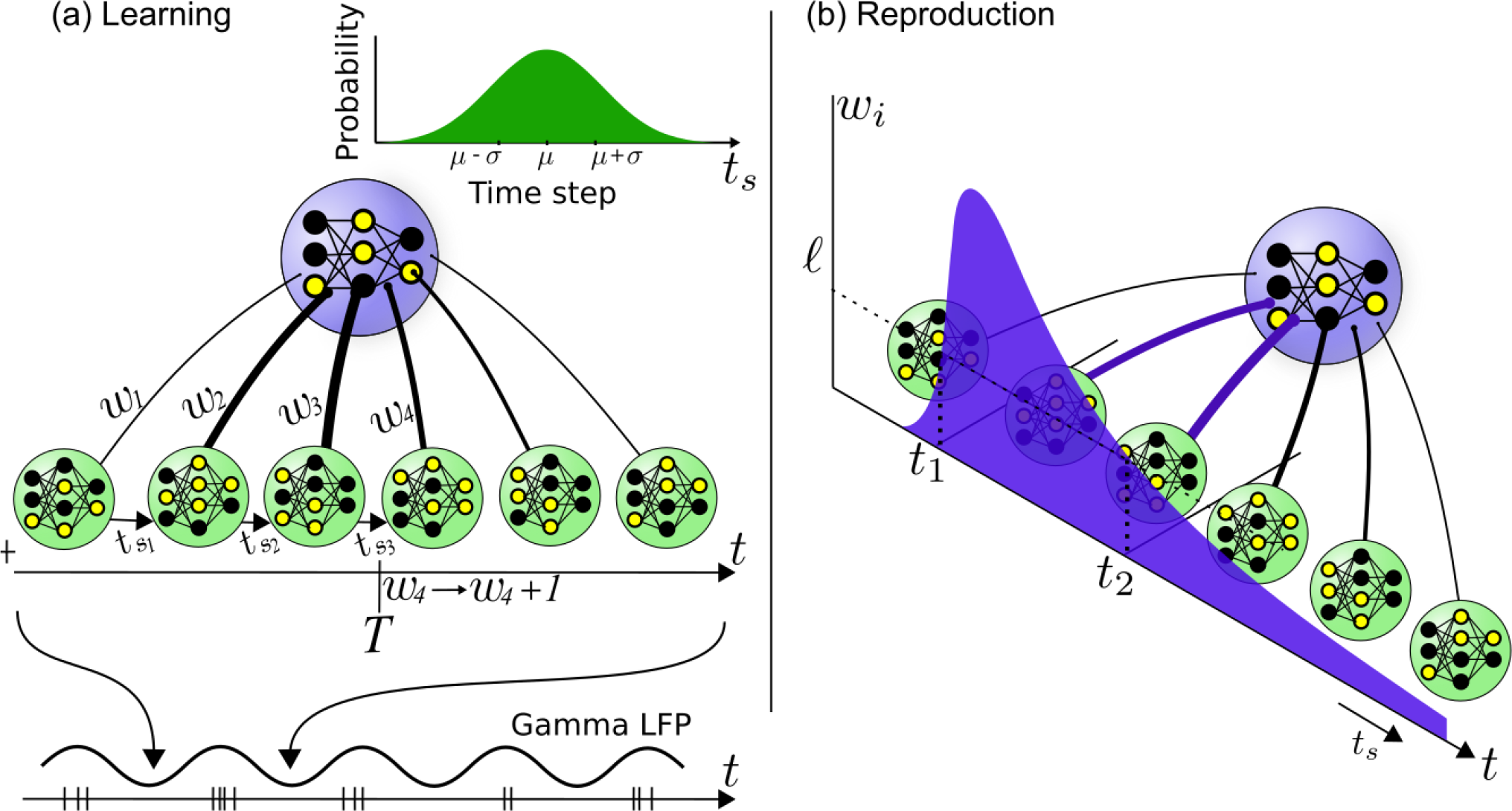
(a) Visual representation of the Learning Phase of the model. Each green sphere represents a state and the lines between the green and the blue spheres represent the weight *w*_*i*_ of the link between the state itself and a specific other neural unit represented by the blue sphere. The time steps *t*_*s*_ compose a Gaussian distribution and the time *t* increases according to these steps. When the time *t* reaches the target time *T*, the weight of current state will raise. (b) Visual representation of the Reproduction Phase of the model. The same states will be activated successively just like the learning phase but linked with a basal response. While the process reaches states with weights above a threshold *ℓ*, it will activate a neural unit responsible to identify the target time.

#### Algorithm 1 Learning phase

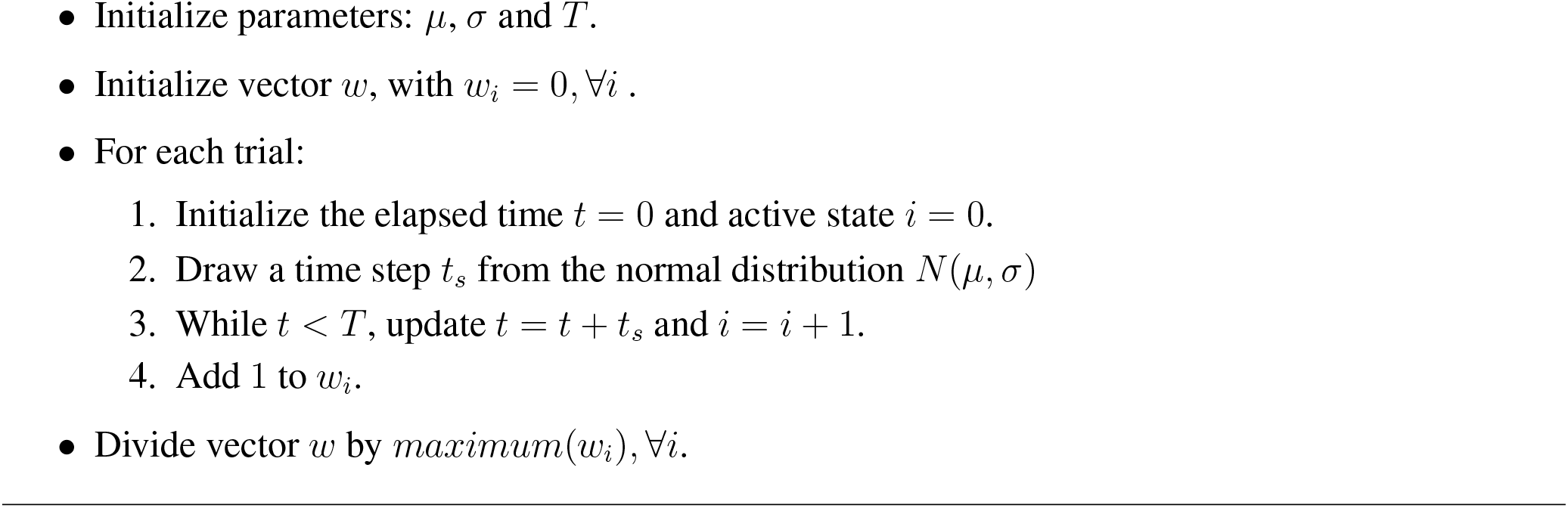

Note that the last activated state *i* after the elapsed time *T* is 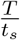. Thus, the function that describes the weight *w*_*i*_ converges to a distribution obtained by a change of variables from the Gaussian distribution of *t*_*s*_ to 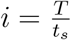. Prior to normalization, we can describe the distribution as:

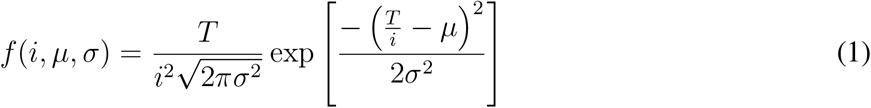

We then reduced the parameters *μ* and *σ* to the coefficient *ρ* = *σ*/*μ*. When we change the parameters of the states *i* to the pressing time using the transformation *t* = *μ* · *i*, and make *σ* = *μρ*, Equation 1 becomes:

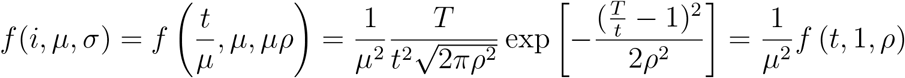

The threshold *ℓ* is the lower degree of activation that generates the behavioral response, and it ranges from 0 to 1. Thus, we have to normalize the maximum weight to 1 (last step in Algorithm 2) to establish a comparison of *w*_*i*_ and *ℓ*. Given the maximum *M* (*ρ*) of the function *f* (*t*, 1, *ρ*), we have that 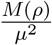 must be the maximum of the function *f*(*i*, *μ*, *σ*). The final function describing the neural activity at the network in terms of the time elapsed is

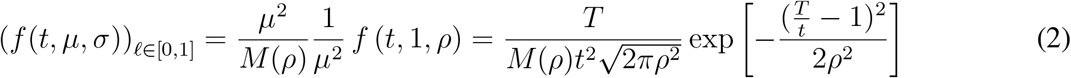

so we can conclude that the only relevant parameters of the model are *ρ* and *ℓ*.

### 2.3 Behavioral Reproduction

In the reproduction phase the model generates single-trial responses that are similar to the responses observed in the experimental peak trials. The model uses the same series of states shown in Figure 2b. When the network activates states whose output weights are lower than a threshold 0 ≤ *ℓ* ≤ 1, rats respond in a basal rate *r*_*b*_. When it reaches states whose weights are larger than the threshold, rats respond in a higher rate *r*_*a*_. Algorithm 2 shows the simulation steps. The rationale behind this scheme is that rats tend to respond in bursts of response around the criterion (Church et al., 1994).

The reproduction phase algorithm generates single-trial responses. We simulated the hypothetical presses using a Poisson distribution with average rate *λ* = *r*_*a*_ between time the burst start *t*_1_ and burst stop *t*_2_, and *λ* = *r*_*b*_ for the remaining period.

### 2.4 Parameters Search

We fitted the parameters to the experimental data of each rat separately. Parameters *ρ* = *σ*/*μ* and *ℓ* are important to the neural mechanism, while *r*_*a*_ and *r*_*b*_ are important to the behavioral response of rats.

Given each of these 4 parameters, we generated trials of the reproduction phase—for each trial, we simulated a set of lever-pressing times and calculated a response-time histogram similar to Figure 1(b) using 80 bins (1 bin per second).

#### Algorithm 2 Reproduction phase

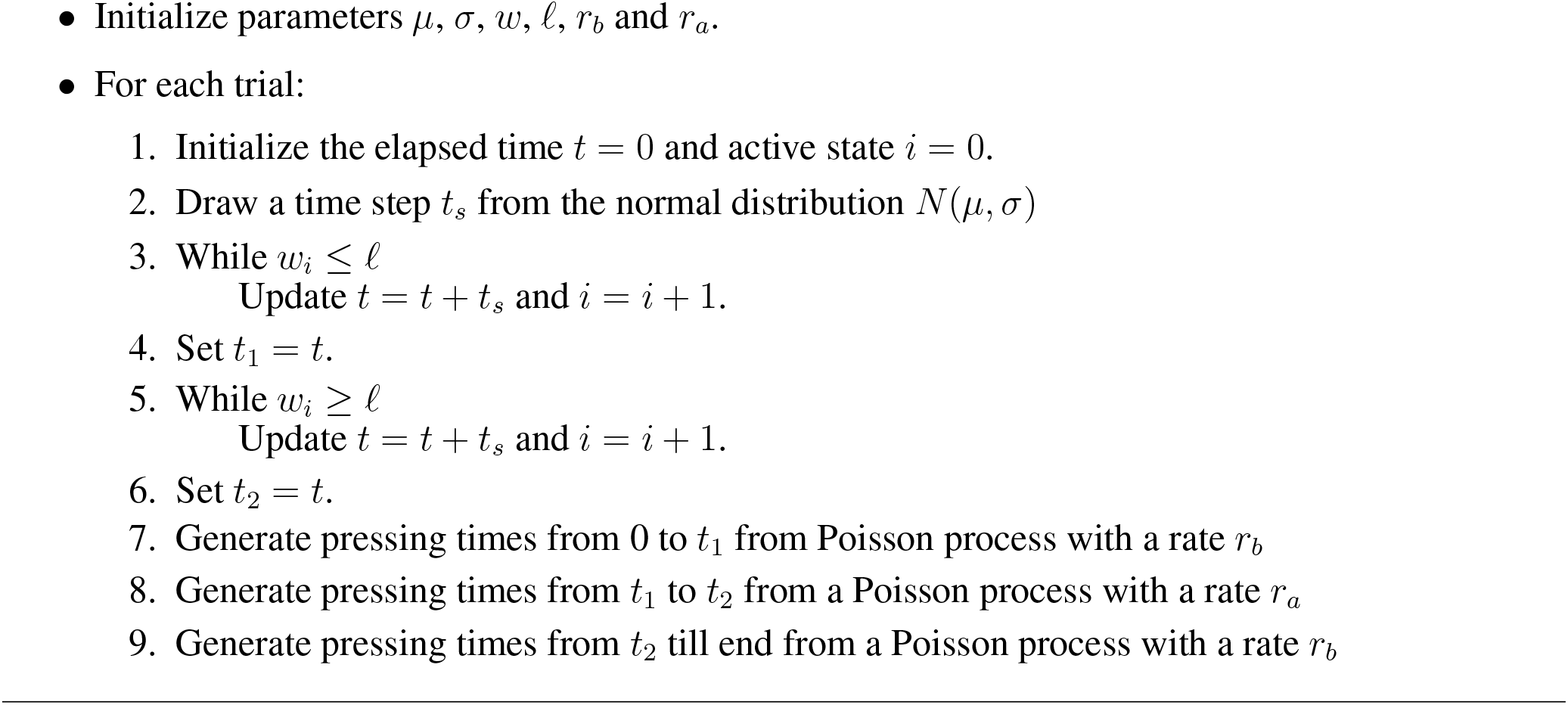

We found the best fit by calculating the Euclidean distance between the model and one specific rat. The difference was between the experimental and the model vector. The experimental vector contained each 1-second bin of the experimental lever press distribution of a rat. The model vector contained each 1-second bin of the distribution generated by the model. After calculating the difference for many possible parameters for the model, we stored the parameters that gave the lower distance.

The basal rate and the burst rate were obtained by analyzing each distribution for each rat. That way, we made sure that the model parameters would be compatible with each rat behavior. The basal rate was acessed from the average pressing rate for times longer than the criterion — the rate at the tail of the experimental distributions from Fig. 2. We searched the other two parameters (*ρ* and *ℓ*) in a grid where both parameters ranged from 0 to 1 in intervals of 0.05.

### 2.5 Weber Law

With the best parameters it is possible to check the validity of the scalar property. To assess the scalar property, we need to change the target time. For each target time, we simulated the model 10 times and averaged the standard deviation of the resulting pressing distributions given by the model. Only 10 times was enough because the distributions had little variation in each simulation.

Lever presses associated during the basal response influence the response distributions to be above the zero line, i.e., the responses distribution do not converge to zero at their tails. Hence, we are not able to normalize these curves and neither calculate their associated statistics. For that reason, we considered only the responses during the bursts to calculate the standard deviations.

### 2.6 Individual trials simulation and analysis

For each rat, we used the parameters search to find the best parameters associated with the model. With these parameters, we can reproduce single trials from the rat by using the Algorithm 2.

To compare the start (*t*_1_) and stop times (*t*_2_) from both experimental data and the model we fitted each experimental trial using the methodology proposed by (Church et al., 1994). In that method, we made a search within *t*_1_ and *t*_2_ and stored the values that gave the maximum value of

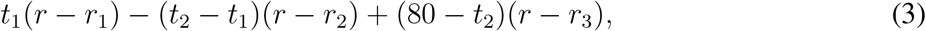

where *r* is the pressing rate for the whole trial, *r*_1_ is the pressing rate for the first interval (from 0 to *t*_1_), *r*_2_ is the pressing rate for the second interval (that is equivalent to the burst rate, from *t*_1_ to *t*_2_) and *r*_3_ is the pressing rate in the last interval (from *t*_2_ to 80 s — the last time bin recorded).

## 3 Results

### 3.1 Reproduction of behavioral responses

The model reproduced the experimental pressing rate response of 6 individual rats (Figure 3a). We characterized the pressing rate by tuning 4 parameters: the *ρ* = *σ*/*μ* ratio, the threshold *ℓ* of the neural network, the burst rate *r*_*a*_, and the basal rate *r*_*b*_. We selected the parameters by minimizing the Euclidean distance (Table 1) between the simulated and experimental responses. Except for rat 3, all simulations could reproduce the target time and the standard deviation of the curve.

**Figure 3:**
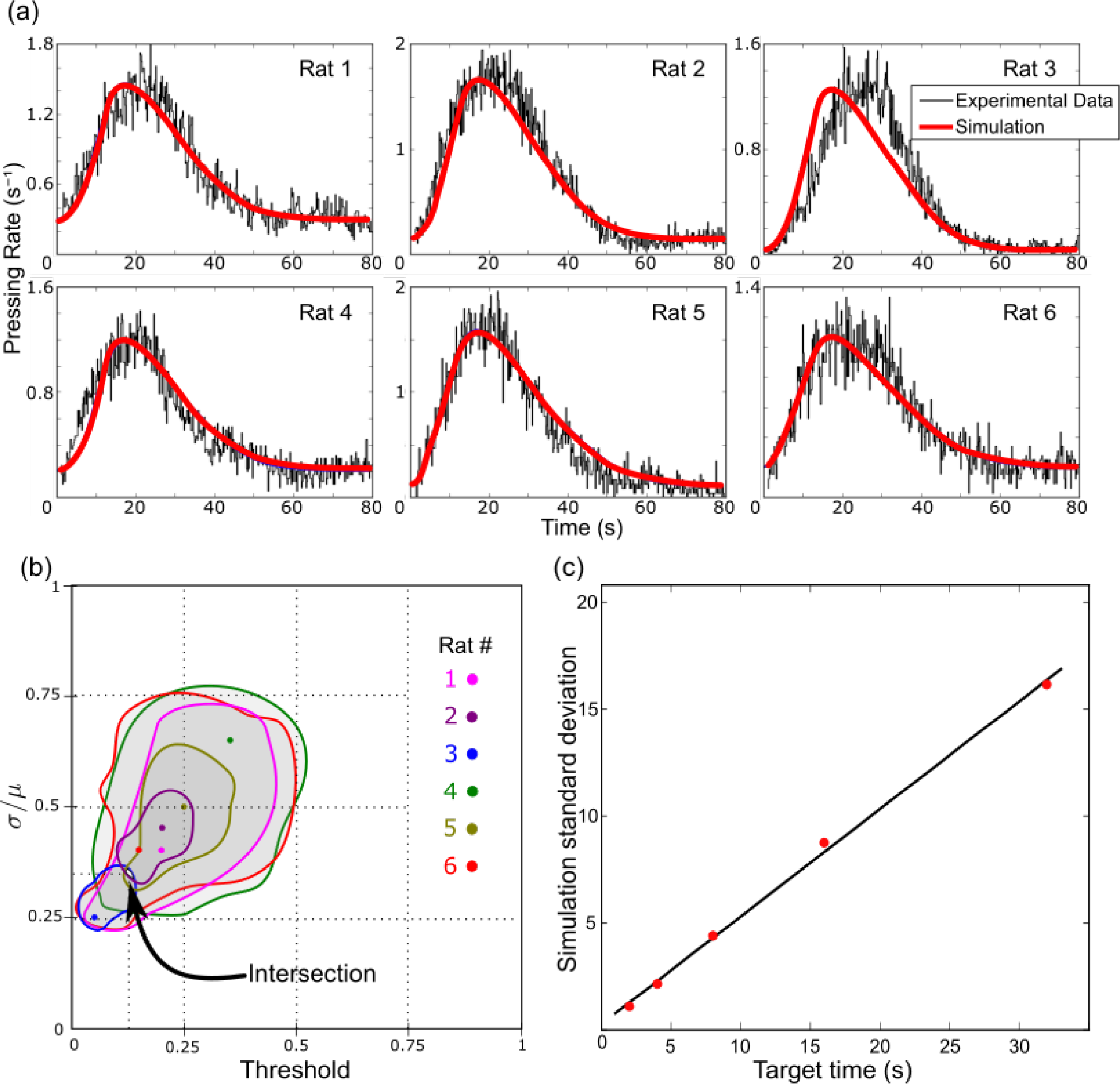
(a) Fitting of the model with the experimental data for each of the 6 rats. The red line represents the simulation fitted with the best parameters and the black line represents the experimental data. (b) Region in the space of parameters in which the goodness of fit measured by the Euclidean Distance between the simulation and the actual data was smaller than 1.2. Each color represents a different rat. The solid dots represent the minimum distance (best goodness of fit). (c) Reproduction of the scalar property by calculating the standard deviation for different target times and then plotting a linear regression.

Each rat had a different pressing rate. We considered in the model that the rat would press at the burst rate *r*_*a*_ when the downstream neural network is active. Otherwise, it would press at the basal rate *r*_*b*_. The target time *T* is the most important factor to determinate the activation of the downstream network. The threshold *ℓ* and the coefficient *ρ* = *σ*/*μ* of the time step sizes dictated the deviation of the final distribution. They are responsible for a fine-tuning of the response curve.

**Table 1:** Euclidean distance between the experimental data and the best fitted model for each rat

| Rat | 1 | 2 | 3 | 4 | 5 | 6 |
| --- | --- | --- | --- | --- | --- | --- |
| Distance (resps./s) | 0.86 | 1.06 | 1.14 | 0.58 | 0.91 | 0.64 |

Notice that the rat 3 had the highest Euclidean distance. That difference happened because the peak of the response distribution was delayed in relation to the peak predicted by the model — the rat was biased.

### 3.2 Parameters similarities

The parameter *ρ* = *σ*/*μ* and *ℓ* obtained for each of the rats were similar. Within the space of parameters, we plotted the regions where the Euclidean distance of the model and the experimental data were less than a certain value (Figure 3b). The smallest Euclidean distance that we found an intersecting region is approximately 1.2. The parameters correspondent to that distance are (*ℓ*, *ρ*) = (0.125, 0.35).

The basal rate found for each rat was, respectively, 0.3, 0.15, 0.02, 0.2, 0.1 and 0.19. The burst rate used for each rat was, respectively, 1.7, 2.1, 1.3, 1.8, 2.2, 1.3. Notice that basal rates were similar between rats, as were burst rates. The basal rates correspond to the height of the pressing rate curves at the immediate beginning and during the end of the sessions.

### 3.3 Weber Law

There was a linear relationship between the target time and the standard deviation of the responses given by the model (Figure 3c). The error within the standard deviations (y-axis) was small and thus not visible in the graph. The Weber fraction *WF* calculated using the parameters given by the intersecting region of the six rats was *WF* = 0.5. The value is consistent with values found in the literature, such as the range of 0.35 to 0.5 in (Gibbon, 1977).

### 3.4 Start and stop time distributions

The distributions of start and stop times that the model generated were similar to the experimental distributions (Figures 4a,b). We simulated single trials using Algorithm 2 for each rat. The parameters used were those from the best fit. The number of trials simulated was proportional to the experimental number of trials for each rat. The model reproduced the peak positions. It could not reproduce well the right tail of the start bursts and left tail of stop bursts. Figures 4c and 4d compare model-generated and experimental data, respectively, from one rat.

**Figure 4:**
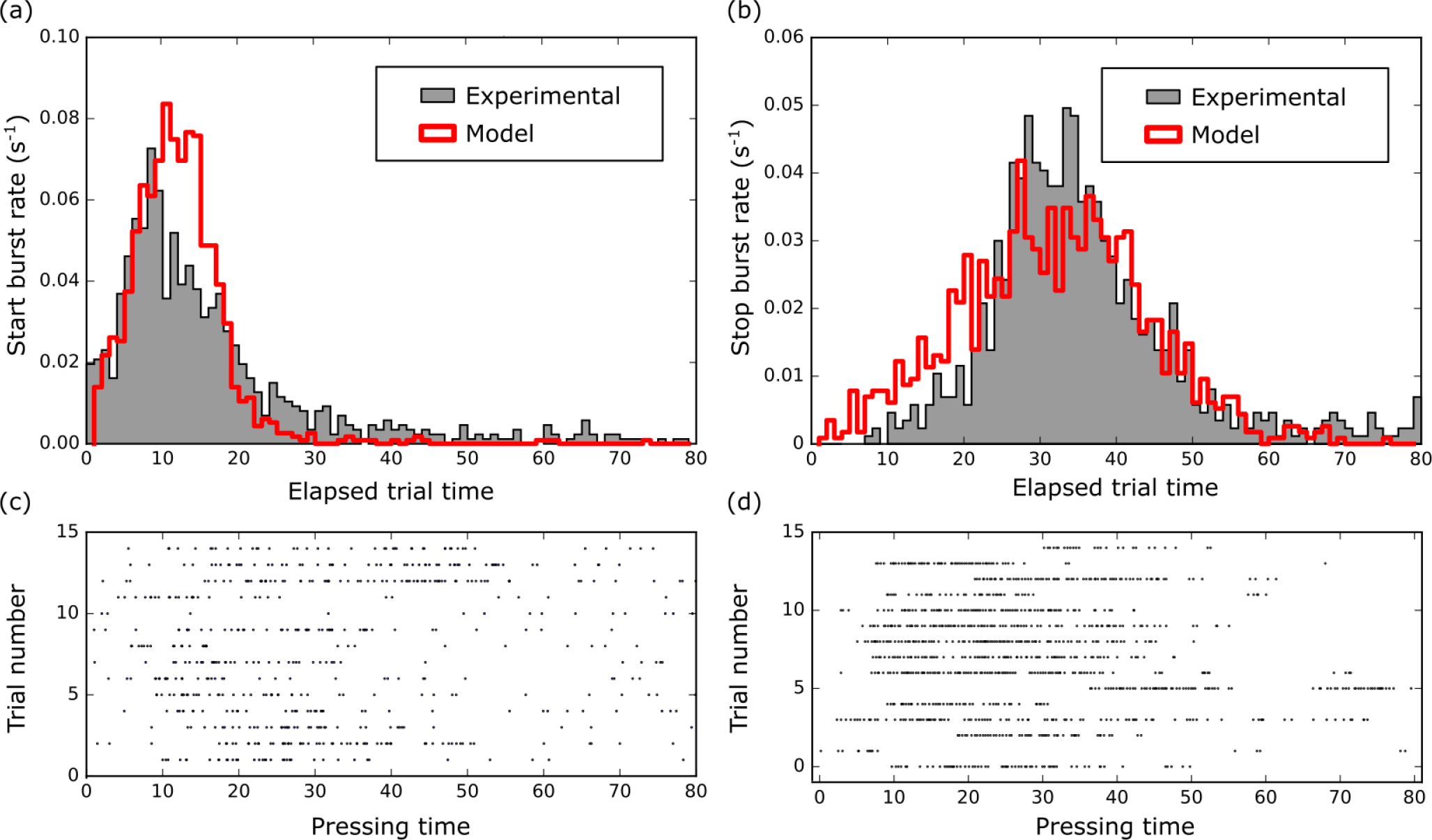
(a) Comparison between start-times distribution in individual trials, both from experimental data (gray) and generated by the model (red). (b) Same as (a) for stop-times. (c) Individual trials generated by the model using the parameters optimized from rat 2. (d) Individual trials for rat 2, session 11.

## 4 Discussion

The proposed model works with neural networks that generate reproducible sequences of states at steady rates. Heteroassociative networks (Sompolinsky and Kanter, 1986; Camargo et al., 2018) can be used to implement our model as a biological neural network, as well as Synfire chains (Abeles, 1991; Miyata et al., 2013). The later introduced the idea of a rapid sequential activation of groups of neurons, representing the states in our model. Accordingly, heteroassociative networks are promising as a biological implementation of the model, since they can produce long sequences of state activation within a single network while performing pattern completion (Camargo et al., 2018). This kind of networks are present in both the CA3 (Lisman et al., 2005) and CA1 (Miyata et al., 2013) subregions of the hippocampus, which is involved in some time-related tasks (Meck et al., 1984). Accumulated evidence suggests that timing mechanisms are decentralized (Paton and Buonomano, 2018) and the hippocampus is one from brain areas where sequences of states could be evoked (Buzsáki and Tingley, 2018).

Neural activity oscillations are the main candidate for providing the steady state transition rates. Subsequent states could be represented on consecutive oscillatory cycles. The parameter *μ* in our model would be the oscillation period and *σ* its variability. Interestingly, the coefficient *ρ* = *σ*/*μ* of the oscillation - not the Weber fraction - could be important to characterize brain oscillations, for instance, indicating its adaptability in certain contexts (Schlee et al., 2014; Calomeni et al., 2017). The parameters obtained from the intersection of the 6 rats (Figure 3b) predicted a coefficient *ρ* at approximately 0.35. This could correspond, for instance, to *μ* = 25*ms* and *σ* = 8.75*ms*, resulting in a frequency range of 30-60Hz, in the range of experimentally observed slow gamma waves (Belluscio et al., 2012). Gamma frequency waves are present in several brain areas and could be a candidate for providing the state transition rates.

The reproducible sequences of states at steady rates would also give rise to cells that respond at specific times of the task, similarly to time cells, discovered in the Hippocampus (Einchenbaum, 2013), medial entorhinal cortex (MEC) (Tsao et al., 2018) and medial prefrontal cortex (mPFC) (Tiganj et al., 2017). Neurons with behavior similar to time cells could appear both as part of the states in the sequence and as part of the downstream network. These time cells would also “retime” when an important temporal parameter changes. (MacDonald et al., 2011). The existence of neurons with time cells is an important link of the proposed model with existing biological evidence.

The proposed model uses a bottom-up approach, from properties of brain neural networks to behavioral responses from rats. Interestingly, it resembles the computational implementation of the Learning-to-Time (LeT) behavioral model. In both models, serially activated states have different link strengths to an outside unit, and a threshold leading to behavioral responses. There are differences, such as the LeT extinguishing of non-active link strengths. The LeT model was motivated by the proposal of behavioral internal states, without reference to the underlying biological mechanisms. Our proposed model can provide a link from the behavioral model to a possible biological substrate.

Recently, a new model based on time cells was proposed (Zeki and Balcı, 2019). The model is close to our peak-interval model in that it is also a possible biologically plausible implementation of the LeT model and can explain the scalar property. Both models are based on a chain of activation, but the time cells model is a chain of single neurons while our model is a chain of neural ensembles. Since the time cell model was implemented using an integrate-and-fire model, to achieve the scalar property they assumed biophysically-plausible inhibitory currents to modulate the interspike intervals.

Our model has some limitations. One example is that the model could not reproduce perfectly the start and the stop distributions of burst press activities (Figure 4). The genesis of this discrepancy may rely on the way we generated single trials. A visual inspection of Figure 4c and d shows that the experimental data is more sparsely distributed than the simulation. That difference comes from the fact that the neural network we are proposing deals with the task-related interval representation in the brain. The way the behavior appears will have more intricate mechanisms. For example, the hypothesis of one single burst per trial is false in all experimental responses — processes like attention, impulsivity, satiation certainly imposes greater level of complexity to behavior that stretches beyond the scope of our model.

Another limitation is that there is not a source of bias in our model. Bias could arise from impulsive responding (Matell and Portugal, 2007; Renda et al., 2014), for example. The model does not capture correctly the pressing rate distribution of rat 3, whose peak response lies 5 to 10 seconds delayed from the target time. The peak of the distribution of simulated responses depends on the target time learned by the rat. One way to account for a variation in the distribution peak would be the addition of a bias. That bias would change the encoding of the neural network to the response rate, and thus change the time between two neural states in the behavioral reproduction phase. Even so, since the bias is more related to the decision-making process than the timing mechanism itself, we decided to not include this parameter and thus reduce the complexity of the model.

## 6 Acknowledgements

During the project, TTV was supported by the Fundação de Amparo à Pesquisa do Estado de São Paulo (FAPESP, grant #2016/19691-1) and by the Universidade de São Paulo International Cooperation Office Scholarship. MSC is a member of the Instituto Nacional de Ciência e Tecnologia sobre Comportamento, Cognição e Ensino, with support from the Brazilian National Research Council (CNPq, Grant #465686/2014-1), the Coordination of Superior Level Staff Improvement (CAPES, Grant #88887.136407/2017-00), and FAPESP (Grant #2014/50909-8). The authors would like to thank the members of the Timing and Cognition Lab (http://neuro.ufabc.edu.br/timing/), at Universidade Federal do ABC, for their suggestions during project development. Finally, authors thank Prof. Armando Machado for helpful discussions during the research.

